# Topographic gradients of intrinsic dynamics across neocortex

**DOI:** 10.1101/2020.07.03.186916

**Authors:** Golia Shafiei, Ross D. Markello, Reinder Vos de Wael, Boris C. Bernhardt, Ben D. Fulcher, Bratislav Misic

## Abstract

The intrinsic dynamics of neuronal populations are shaped by both macroscale connectome architecture and microscale attributes. Neural activity arising from the interplay of these local and global factors therefore varies from moment to moment, with rich temporal patterns. Here we comprehensively characterize intrinsic dynamics throughout the human brain. Applying massive temporal feature extraction to regional haemodynamic activity, we estimate over 6,000 statistical properties of individual brain regions’ time series across the neocortex. We identify two robust topographic gradients of intrinsic dynamics, one spanning a ventromedial-dorsolateral axis and the other spanning a unimodal-transmodal axis. These gradients are distinct in terms of their temporal composition and reflect spatial patterns of microarray gene expression, intracortical myelin and cortical thickness, as well as structural and functional network embedding. Importantly, these gradients are closely correlated with patterns of functional activation, differentiating cognitive *versus* affective processing and sensory *versus* higher-order cognitive processing. Altogether, these findings demonstrate a link between microscale and macroscale architecture, intrinsic dynamics, and cognition.

## INTRODUCTION

The brain is a complex network of anatomically connected and perpetually interacting neuronal populations [87]. Inter-regional connectivity promotes signaling via electrical impulses, generating patterned electrophysiological and haemodynamic activity [6, 89]. Neuronal populations are organized into a hierarchy of increasingly polyfunctional neural circuits [47, 53, 65], manifesting as topographic gradients of molecular and cellular properties that smoothly vary between unimodal and transmodal cortices [50]. Recent studies have demonstrated cortical gradients of gene transcription [12, 31], intracortical myelin [49], cortical thickness [100] and laminar profiles [72].

The topological and physical embedding of neural circuits in macroscale networks and microscale gradients influence their dynamics [40, 56, 102]. For a neuronal population, the confluence of local properties and global connectivity shapes both the generation of local rhythms, as well as the propensity to communicate with other populations. Numerous studies, mainly using electrophysiological recordings, suggest that intrinsic timescales systematically vary over the cortex [32, 51, 61, 71]. The primary functional consequence of this hierarchy of timescales is thought to be a hierarchy of temporal receptive windows: time windows in which a newly arriving stimulus will modify processing of previously presented information [7, 15, 16, 43, 48, 50]. Thus, areas at the bottom of the hierarchy preferentially respond to immediate changes in the sensory environment, while areas at the top of the hierarchy preferentially respond to more long-lived or contextual information.

The relationship between structure and dynamics is also observed at the network level [89]. Intrinsic or “resting state” networks possess unique spectral fingerprints [54]. Signal variability, measured in terms of standard deviations or entropy, is closely related to structural and functional network embedding [14, 33, 69, 84]. More generally, the autocorrelation properties of blood oxygenation level-dependent (BOLD) signal are correlated with topological characteristics of structural brain networks, such that areas with greater connectivity generate signals with greater autocorrelation [25, 82]. Finally, in nonlinear dynamic models, highly interconnected hubs exhibit slower dynamic fluctuations, while sensory areas exhibit fast fluctuating neural activity [40]. Indeed, nonlinear dynamic models offer better fits to empirical functional connectivity if they assume heterogeneous local dynamics that follow a unimodal–transmodal gradient [17, 22, 101].

Altogether, multiple lines of evidence suggest that local computations may reflect systematic variation in microscale properties and macroscale network embedding, manifesting as diverse temporal properties of regional activity. However, conventional computational analysis is based on specific, manually selected temporal features (autocorrelation, variance, spectral power) and specific anatomical properties (gene expression, cortical thickness, connectivity). Yet the time series analysis literature is vast and interdisciplinary; how do other metrics of temporal structure vary across the brain and what can they tell us about cortical organization? Do different types of local computations manifest as different organizational gradients? Here we comprehensively chart intrinsic dynamics across the cerebral cortex, mapping temporal organization to structural organization. We apply massive temporal feature extraction to resting state BOLD signals to derive an exhaustive intrinsic dynamic profile for each brain region. We then systematically investigate the relationship between local temporal properties and gene expression, microstructure, morphology, structural connectivity and functional connectivity. We show that intrinsic dynamics reflect molecular and cytoarchitectonic gradients, as well as patterns of structural and functional connectivity.

## RESULTS

All analyses were performed on four resting state fMRI runs from the Human Connectome Project [93]. The data were pseudorandomly divided into two samples of unrelated participants to form *Discovery* and *Validation* samples with *n* = 201 and *n* = 127, respectively [21]. External replication was then performed using data from the Midnight Scan Club [41]. Massive temporal feature extraction was performed using highly comparative timeseries analysis, *hctsa* [29, 30], yielding approximately 6,000 features per regional time series, including measures of frequency composition, variance, autocorrelation, fractal scaling and entropy (Fig. 1). The results are organized as follows. We first investigate whether regions that are structurally and functionally onnected display similar intrinsic dynamics. We then characterize the topographic organization of temporal features in relation to microstructural attributes and cognitive ontologies.

**Figure 1.**
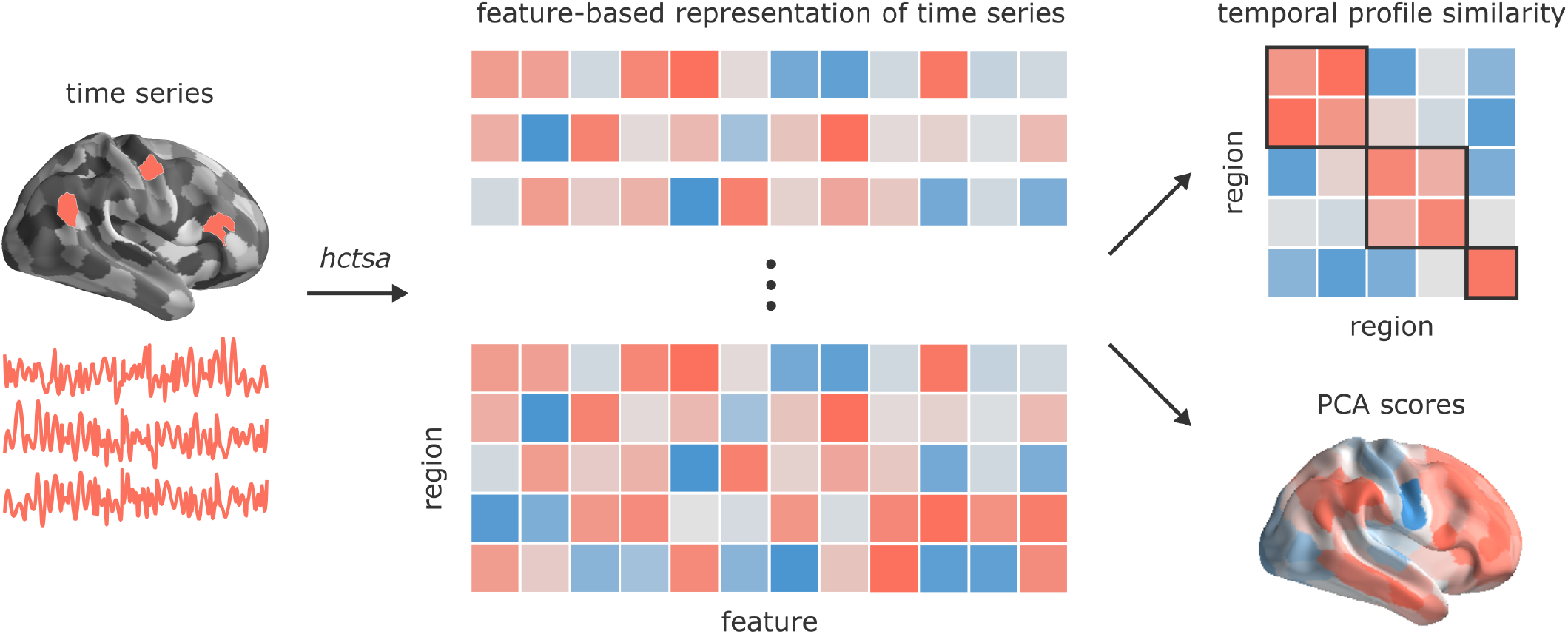
Temporal phenotyping of regional dynamics. The highly comparative time-series analysis toolbox, *hctsa* [29, 30], was used to extract over 6,000 temporal features of the parcellated time series for each brain region and participant, including measures of autocorrelation, variance, spectral power, entropy, etc. Regional dynamic profiles were then entered into two types of analyses. In the first analysis, pairs of regional temporal feature vectors were correlated to generate a region × region temporal profile similarity network. In the second analysis, principal component analysis (PCA) was performed to identify orthogonal linear combinations of temporal features that vary maximally across the cortex.

### Inter-regional temporal profile similarity reflects network geometry and topology

We first assessed the extent to which intrinsic dynamics depend on inter-regional physical distance, anatomical connectivity and functional connectivity. We estimated similarity between inter-regional dynamics by computing Pearson correlation coefficients between regional temporal feature vectors (Fig. 1). Two regional time series are judged to be similar if they have similar temporal profiles, estimated across a comprehensive and diverse set of temporal properties (e.g. similar similar entropy, stationarity, linear correlation properties) [27]. This measure of similarity identifies pairs of regions that have similar dynamical features, but not necessarily coherent or synchronous dynamics (Fig. 2a). We refer to correlations between regional temporal feature profiles as “temporal profile similarity”.

**Figure 2.**
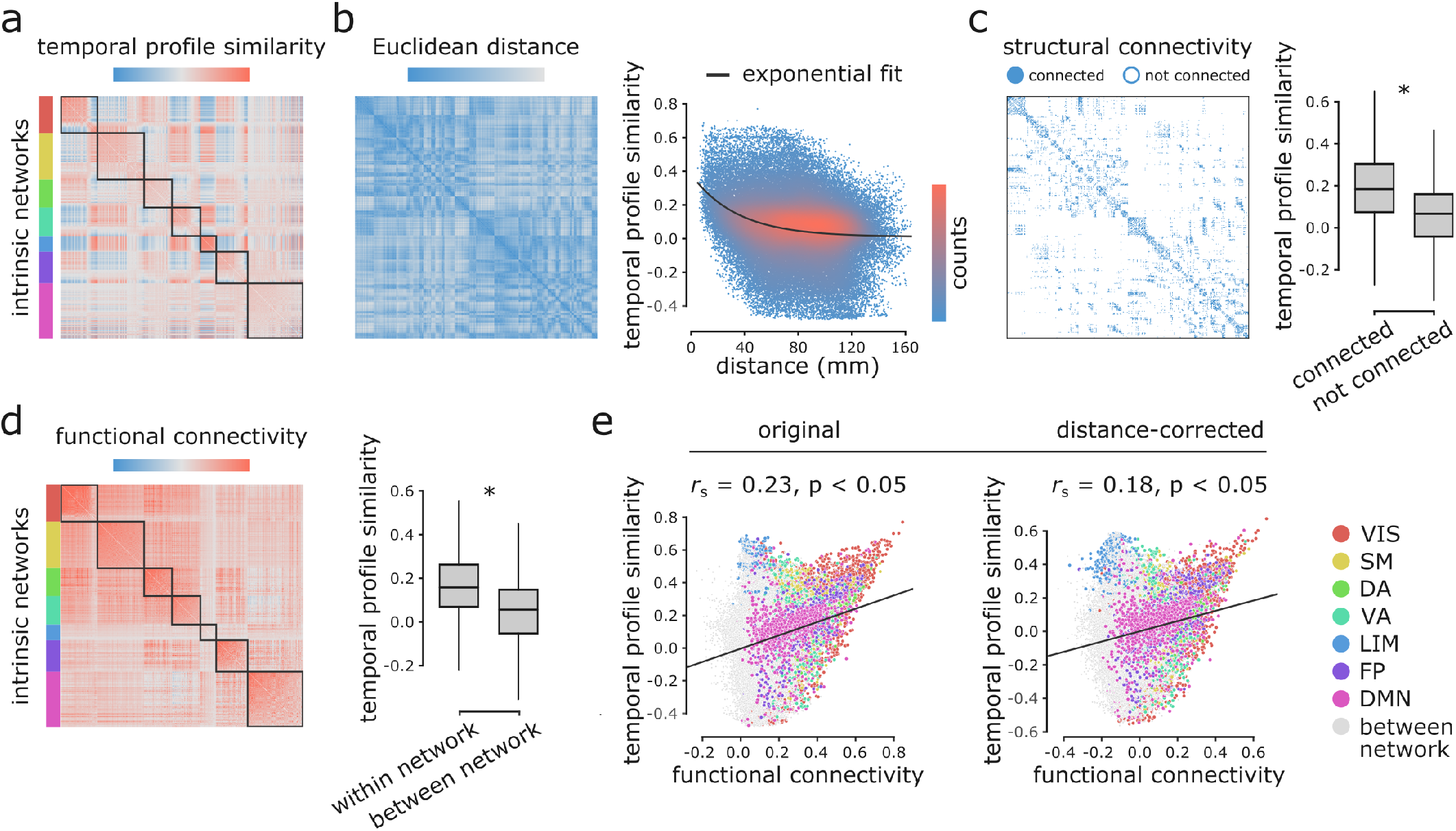
Inter-regional temporal profile similarity reflects network geometry and topology. (a) Temporal profile similarity networks are constructed by correlating pairs of regional temporal feature vectors. Brain regions are ordered based on their intrinsic functional network assignments [106]. (b) Temporal profile similarity between regions significantly decreases as a function of Euclidean distance between them. The black line represents an exponential fit as *y* = 0.37*e*^-0.03*x*^ + 0.01, where *y* is temporal profile similarity and *x* is Euclidean distance. (c, d) Regional temporal features are compared between pairs of cortical areas using their structural and functional connectivity profiles. Pairwise temporal profile similarity is significantly higher among structurally-connected areas (c), and among regions that belong to the same intrinsic functional networks (d). Statistical significance of the observations is indicated using an asterisk (two-tailed t-test; *P* ≈ 0). (e) Temporal profile similarity is positively correlated with functional connectivity. This relationship remains after partialling out Euclidean distance between regions from both measures using exponential trends. *r*_s_ demonstrates Spearman rank correlation coefficient. Linear regression lines are added to the scatter plots for visualization purposes only. Connections are colour-coded based on the Yeo intrinsic networks [106]. VIS = visual, SM = somatomotor, DA = dorsal attention, VA = ventral attention, LIM = limbic, FP = fronto-parietal, DMN = default mode.

Fig. 2b shows a negative exponential relationship between spatial proximity and temporal profile similarity, meaning that regions that are spatially close exhibit similar intrinsic dynamics. Interestingly, regions that share an anatomical projection have greater temporal profile similarity than those that do not (Fig. 2c; two-tailed *t*-test; *P* ≈ 0). To test whether this anatomically-mediated similarity of dynamical features is not due to spatial proximity, we performed two additional comparisons. First, we regressed out the exponential trend identified above, and repeated the analysis on the residuals, yielding a significant difference in temporal profile similarity between connected and non-connected regions (two-tailed t-test; P ≈ 0). Second, we generated an ensemble of 10 000 degree- and edge length-preserving surrogate networks [9], and compared the difference of the means between connected and non-connected pairs in the empirical and surrogate networks. Again, we observe a significant difference in temporal profile similarity between connected and non-connected regions (two-tailed; P = 0.0001).

Likewise, regions belonging to the same intrinsic functional network have greater temporal profile similarity compared to regions in different networks (Fig. 2d; twotailed *t*-test; *P* ≈ 0). We confirmed this finding is not driven by spatial proximity, we repeated the analysis with distance-residualized values, finding a significant difference (two-tailed *t*-test; *P* ≈ 0). We also repeated the analysis using a nonparametric label-permutation null model with preserved spatial autocorrelation [2], again finding significantly greater with-compared to between-network temporal profile similarity (two-tailed; *P*_spin_ = 0.006). More generally, we find a weak positive correlation between temporal profile similarity and functional connectivity (original: Spearman rank *r*_s_ = 0.23, P ≈ 0; distance-corrected: *r*_s_ = 0.18, *P* ≈ 0; Fig. 2e), suggesting that areas with similar temporal features exhibit coherent spontaneous fluctuations, but that the two are only weakly correlated. Fig. 2e shows the correlation between the two; points represent pairwise relationships and are coloured by their membership in intrinsic networks [106]. In other words, two regions could display similar time series features, suggesting common function, but they do not necessarily fluctuate coherently. Thus, representing time series using sets of features provides a fundamentally different perspective compared to representing them as the raw set of ordered BOLD measurements. Altogether, we find that the organization of intrinsic dynamics is closely related to both the geometric and topological embedding of brain regions in macroscale networks.

### Intrinsic dynamics reflect microscale and macroscale hierarchies

We next investigate the topographic organization of temporal features. Applying principal component analysis (PCA) to the region × feature matrix yielded mutually orthogonal patterns of intrinsic dynamics (Fig. 1), with the top two components collectively accounting for more than 70% of the variance in temporal features (Fig. 3a). Fig. 3a shows the spatial distribution of the top two components. The first component (PC1) mainly captures differential intrinsic dynamics along a ventromedial-dorsolateral gradient, separating occipital-parietal cortex and anterior temporal cortex. The second component (PC2) captures a unimodal-transmodal gradient, reminiscent of recently reported miscrostruc-tural and functional gradients [50]. Both components show considerable hemispheric symmetry. In the following sections, we focus on these two components because of their (a) effect size (percent variance accounted for), (b) close resemblance to previously reported topographic gradients, and (c) reproducibility (only the first two components were reproducible in both the HCP and MSC datasets; see *Sensitivity and replication analyses* below). Note that neither spatial maps were significantly correlated with temporal signal-to-noise ratio map, computed as the ratio of the time series mean to standard deviation (tSNR; PC1: *r*_s_ = 0.28, *P*_spin_ = 0.19; PC2: *r*_s_ = 0.21, *P*_spin_ = 0.16).

**Figure 3.**
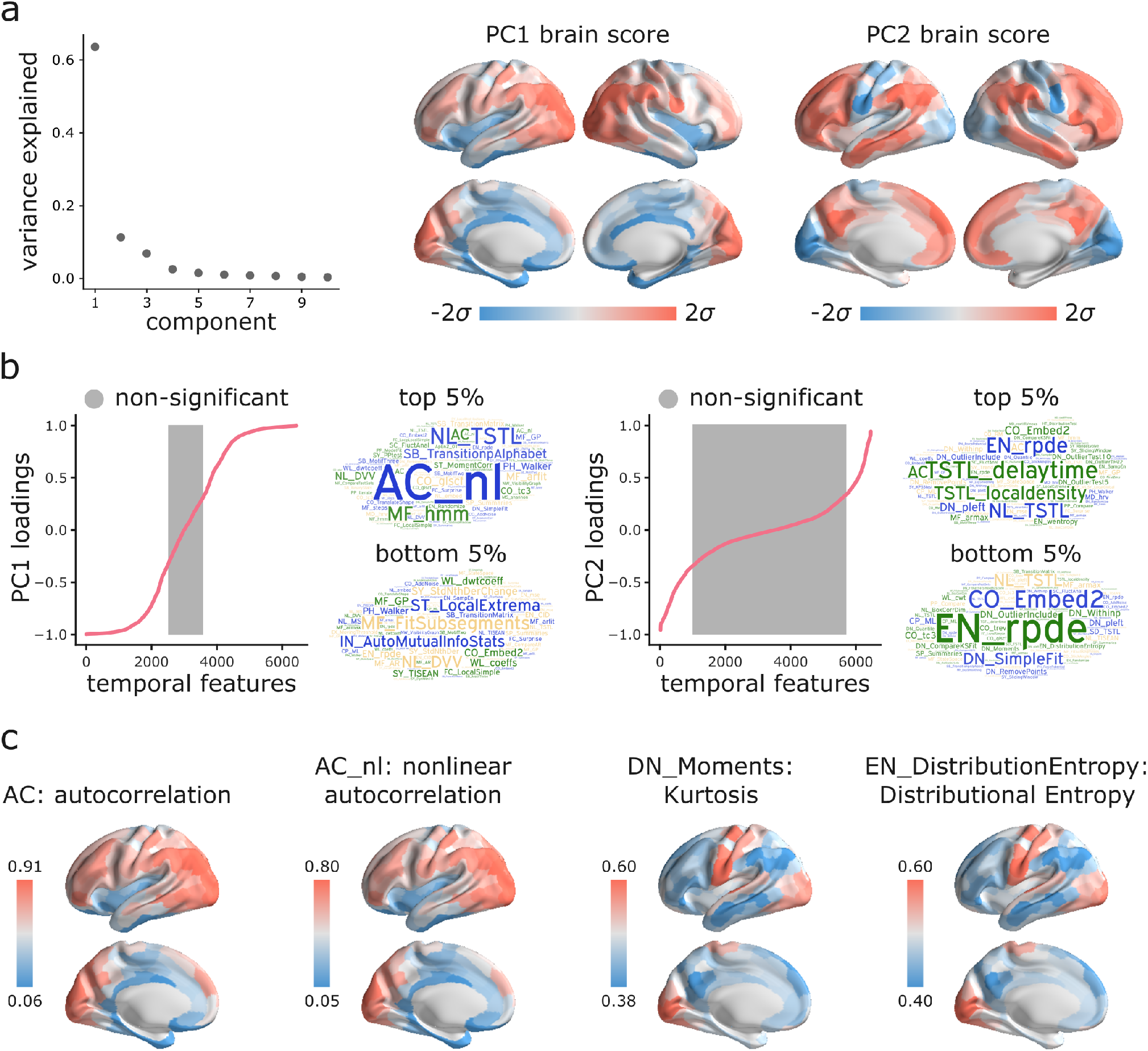
Topographic gradients of intrinsic dynamics. (a) PCA analysis identified linear combinations of hctsa temporal features with maximum variance across the human cortex. Collectively, the first two components (PC1 and PC2) account for 75% of the total variance in time-series features of BOLD dynamics across brain regions. To estimate the extent to which cortical regions display the patterns of intrinsic dynamics captured by each component, *hctsa* matrices were projected back onto the PC weights (eigenvectors), yielding spatial maps of brain scores for each component. Spatial maps are depicted based on the standard deviation *σ* of their respective brain score distributions. (b) To understand the feature composition of the intrinsic dynamic patterns captured by PC1 and PC2, feature loadings were computed by correlating individual *hctsa* feature vectors with the PC score maps. PC loadings thus estimate the shared spatial variance between an individual temporal feature and the composite intrinsic dynamic map captured by a PC. Temporal features are ordered by their individual loadings. Grey indicates non-significance based on 10,000 spatial permutation tests (FDR correction, *α* = 0.001). Features corresponding to the top and bottom 5% of PC1 and PC2 are visualized using word clouds. The complete list of features ranked by loading, their definitions, correlations and *P*-values for both components is presented in machine-readable format in Supporting Tables S2,3. Feature nomenclature in *hctsa* is organized such that the term prefix indicates the broad class of measures (e.g. AC = autocorrelation, DN = distribution) and the term suffix indicates the specific measure (for a complete list, see https://hctsa-users.gitbook.io/hctsa-manual/list-of-included-code-files). (c) The spatial distributions of two high-loading representative time-series features for each component, including AC_1 (first-order linear autocorrelation), AC_nl (nonlinear autocorrelation), DN_Moments_4 (fourth moment or kurtosis of the time-series points distribution), EN_DistributionEntropy (entropy of the time-series points distribution).

Which temporal properties contribute most to these topographic gradients of intrinsic dynamics? To address this question, we systematically assess the feature composition of PC1 and PC2. We compute univariate correlations (i.e. loadings) between individual temporal feature vectors and PC scores (Fig. 3b). Each loading is assessed against 10,000 spin tests and the results are corrected for multiple comparisons by controlling the false discovery rate (FDR; *α* = 0.001; [8]). The top 5% positively and negatively correlated features are shown in word clouds. The complete list of features ranked by loading, their definitions, loadings and *P*-values for both components is presented in machine-readable format in Supporting Tables S2,3. For PC1, consistent with previous reports, we observe strong contributions from multiple measures of autocorrelation (e.g., AC_1, first-order linear autocorrelation; AC_nl, nonlinear autocorrelation; IN_AutoMutualInfoStats, automutual information). For PC2, we observe strong contributions from measures of distribution shape, often captured by measures of distributional entropy (e.g., EN_DistributionEntropy_ks, entropy of kernel-smoothed distribution; DN_Moments_4, kurtosis; DN_pleft_05, how the distribution is balanced about the mean). In other words, PC2 appears to index dynamic range, or how much of the probability density is away from the mean. Interestingly, none of the odd moments (distribution asymmetry) are high in the PC2 loading list, just even moments, suggesting that PC2 captures the shape of the deviations of time-series data points in both directions from the mean.

To illustrate the topographic organization of these attributes, Fig. 3c shows the spatial distributions of two high-loading representative time-series features for each component. Ventromedial areas (lower in the PC1 gradient) have lower linear and nonlinear autocorrelation, while doroslateral areas (higher in the PC1 gradient) have greater autocorrelation. Sensory areas (lower in the PC2 gradient) have greater distributional entropy and kurtosis, while transmodal areas (higher in the PC2 gradient) have lower distributional entropy and kurtosis.

To assess whether the dominant variation in temporal properties of BOLD dynamics varies spatially with structural and functional gradients, we next quantify the concordance between PC1/PC2 and other microstructural and functional attributes (Fig. 4). We use Spearman rank correlations throughout, as they do not assume a linear relationship among variables. Given the spatially autocorrelated nature of both *hctsa* features and other imaging features, we assess statistical significance with respect to nonparametric spatial autocorrelation-preserving null models [2]. PC1 topography is correlated with the first principal component of microarray gene expression computed from the Allen Institute Human Brain Atlas [12, 45] (*r*_s_ = 0.57, *P*_spin_ = 0.03), but no other attributes. PC2 topography is significantly correlated with the first principal component of microarray gene expression (*r*_s_ = −0.45, *P*_spin_ = 0.0008), with the principal gradient of functional connectivity estimated using diffusion map embedding [18, 57, 63] (https://github.com/satra/mapalign) (*r*_s_ = 0.77, *P*_spin_ = 0.0001), with intracortical myelin as measured by T1w/T2w ratio (*r*_s_ = −0.57, *P*_spin_ = 0.0001) [49], and with cortical thickness (*r*_s_ = 0.43, *P*_spin_ = 0.008) [100]. Altogether, the two topographic gradients of intrinsic dynamics closely mirror molecular and microstructural gradients, suggesting a link between regional structural properties and regional dynamical properties. Fig. S1 further confirms this intuition, showing the mean score of each component for three well-known cortical partitions, including intrinsic functional networks [106], cytoarchitectonic classes [96–98] and laminar differentiation levels [66].

**Figure 4.**
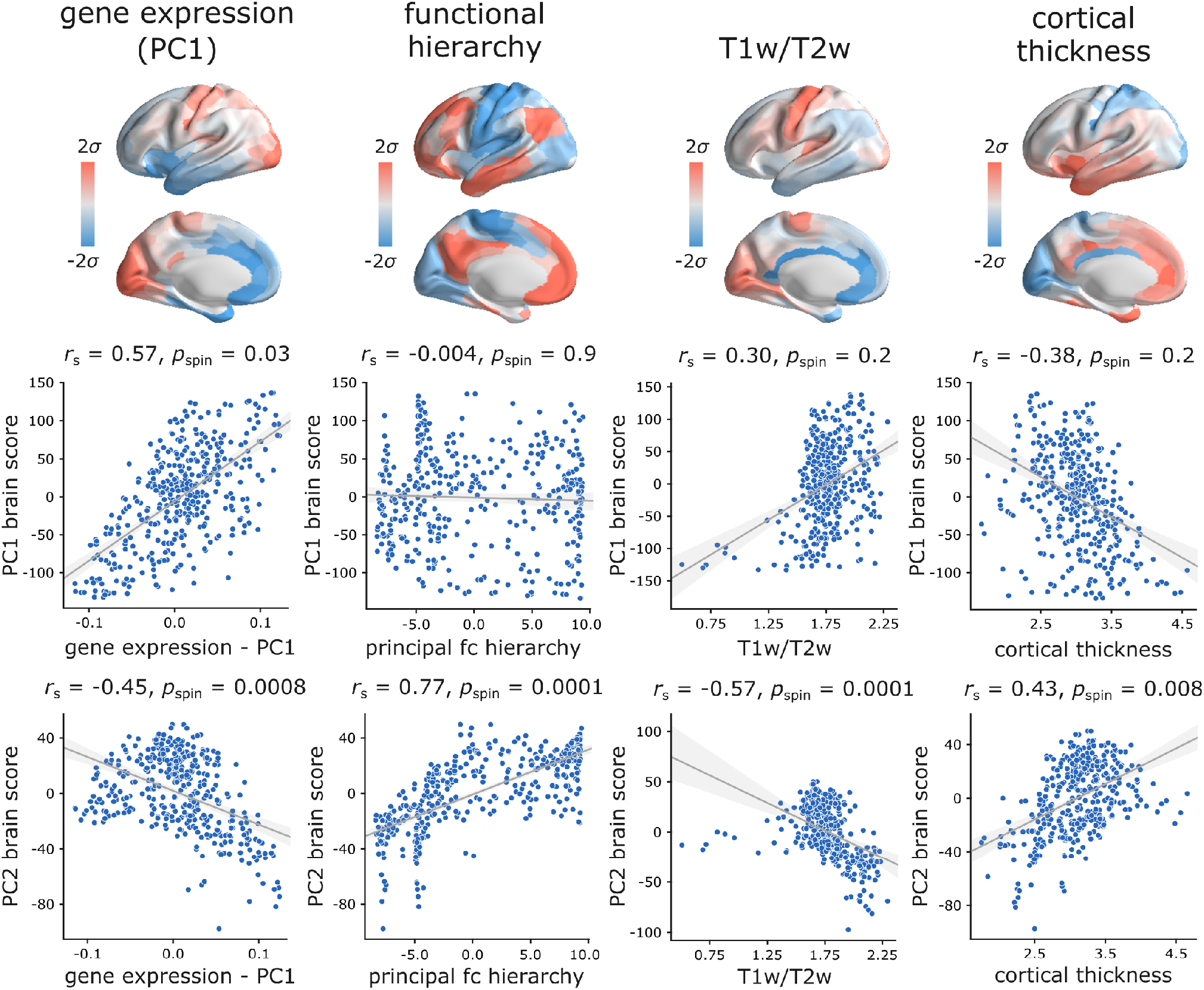
Hierarchical organization of intrinsic dynamics. PC1 and PC2 brain score patterns are compared with four molecular, microstructural and functional maps. These maps include the first principal component of microarray gene expression data from the Allen Human Brain Atlas [12, 45], the first (principal) gradient of functional connectivity estimated using diffusion map embedding [18, 57, 63], group-average intracortical myelin estimated by T1w/T2w ratio, and group-average cortical thickness. The three latter indices were computed from the HCP dataset [93]. Statistical significance of the reported Spearman rank correlation *r*_s_ is assessed using 10,000 spatial permutations tests, preserving the spatial autocorrelation in the data (“spin tests”; [2]). Linear regression lines are added to the scatter plots for visualization purposes only.

### Spatial gradients of intrinsic dynamics support distinct functional activations

Given that topographic patterns of intrinsic dynamics run parallel to distinct microstructural and functional gradients, and marked by distinct temporal properties, we next asked whether these topographic patterns of intrinsic dynamics are related to patterns of functional activation and psychological processes. To address this question, we used Neurosynth to derive probability maps for multiple psychological terms [105]. The term set was restricted to those in the intersection of terms reported in Neurosynth and in the Cognitive Atlas [74], yielding a total of 123 terms (Table S1). Each term map was correlated with the PC1 and PC2 score maps to identify topographic distributions of psychological terms that most closely correspond to patterns of intrinsic dynamics (Bonferroni corrected, *α* = 0.05; Fig. 5). Consistent with the intuition developed from comparisons with intrinsic networks, PC1 intrinsic dynamics mainly defined a cognitive-affective axis (e.g. “attention”, “anticipation” *versus* “stress”, “fear”, “loss”, “emotion”; Fig. 5a), while PC2 dynamics defined a sensory-cognitive axis (e.g. “perception”, “multisensory”, “facial expression” *versus* “cognitive control”, “memory retrieval”, “reasoning”; Fig. 5b).

**Figure 5.**
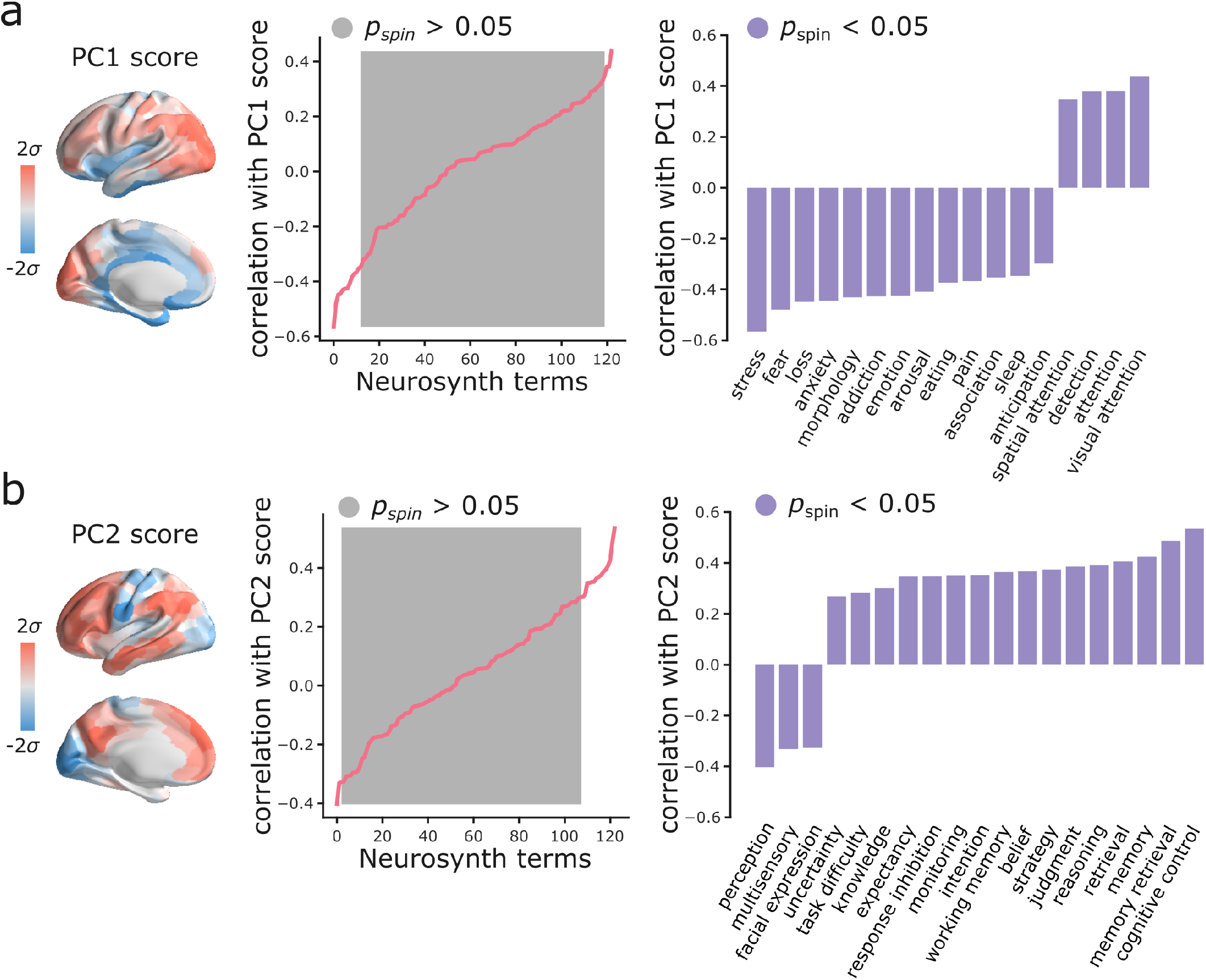
Spatial gradients of intrinsic dynamics support distinct functional activations. We used Neurosynth to derive probability maps for multiple psychological terms [105]. The term set was restricted to those in the intersection of terms reported in Neurosynth and in the Cognitive Atlas [74], yielding a total of 123 terms (Table S1). Each term map was correlated with the PC1 (a) and PC2 (b) score maps to identify topographic distributions of psychological terms that most closely correspond to patterns of intrinsic dynamics. Grey indicates non-significance based on 10,000 spatial permutation tests (Bonferroni correction, *α* = 0.05). Statistically significant terms are shown on the right.

### Sensitivity and replication analyses

As a final step, we sought to assess the extent to which the present findings are replicable under alternative processing choices and in other samples (Fig. 6). For all comparisons, we correlated PC1 and PC2 scores and weights obtained in the original analysis and in each new analysis. Significance was assessed using spatial autocorrelation preserving nulls as before. We first replicated the results in individual subjects in the *Discov-ery* sample by applying PCA to individual region × feature matrices and aligning PCA results through an iterative process using Procrustes rotations (https://github.com/satra/mapalign [57]). The mean individual-level PC scores and weights were then compared to the original findings (Fig. 6a). We next replicated the results by repeating the analysis after grey-matter signal regression (similar to global signal regression as the global signal is shown to be a grey-matter specific signal following sICA+FIX) [36, 37], with near identical results (Fig. 6b). To assess the extent to which results are influenced by choice of parcellation, we repeated the analysis using the 68-region Desikan-Killiany anatomical atlas [23], which were then further divided into 200 approximately equally-sized cortical areas. Again, we find near-identical results (Fig. 6c).

**Figure 6.**
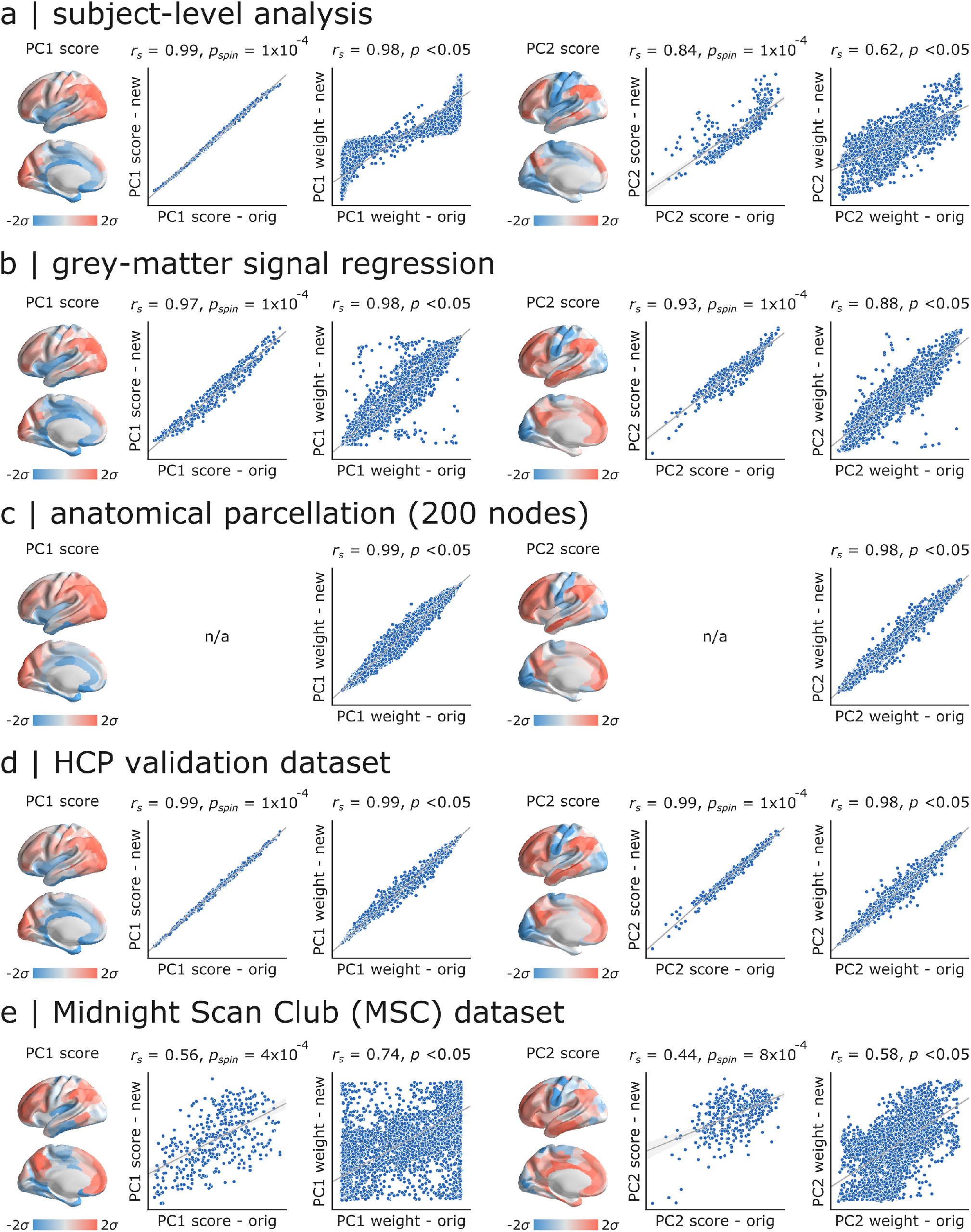
Sensitivity and replication analyses. For all comparisons, we correlated PC1 and PC2 scores and weights obtained in the original analysis and in each new analysis. Significance was assessed using spatial autocorrelation preserving nulls. Specific analyses include: (a) comparing group-level and individual subject-level results, (b) comparing data with and without greymatter signal regression, (c) comparing functional (Schaefer) and anatomical parcellations (Desikan-Killiany), (d) comparing HCP *Discovery* and *Validation* datasets, (e) comparing HCP *Discovery* and Midnight Scan Club datasets.

In the last two analyses we focused on out-of-sample validation. We first repeated the analysis on the held-out *Validation* sample of *n* = 127 unrelated HCP subjects, with similar results (Fig. 6d). Finally, we repeated the analysis using data from the independently collected Midnight Scan Club (MSC) dataset, again finding highly consistent results (Fig. 6e).

## DISCUSSION

In the present report we comprehensively characterize intrinsic dynamics across the cortex, identifying two robust spatio-temporal patterns. The patterns are distinct in terms of their temporal composition and follow microscale gradients and macroscale network architecture. Importantly, the two patterns underlie distinct psychological axes, demonstrating a link between brain architecture, intrinsic dynamics, and cognition.

Our results demonstrate that regional haemodynamic activity, often overlooked in favour of electrophysiological measurements with greater temporal resolution, possesses a rich dynamic signature [35, 58, 60, 75, 92]. While multiple reports have suggested the existence of a timescale or temporal receptive window hierarchy [39, 43, 48, 51, 56, 71, 103?], these investigations typically involved (a) incomplete spatial coverage, making it difficult to quantitatively assess correspondence with other microscale and macroscale maps, and (b) *a priori* measures of interest, such as spectral power or temporal autocorrelation, potentially obscuring other important dynamical features. Although temporal autocorrelation clearly has an important role in the present results, as it does in electrophysiological and electromagnetic recordings, we identify a much broader spectrum of temporal features that relate to microstructure, connectivity and behaviour.

Applying an unbiased, data-driven feature extraction method to high-resolution BOLD fMRI, we decompose intrinsic dynamics into two modes, with distinct topographic organization and temporal properties. One pattern, characterized by variation in signal autocorrelation, follows a ventromedial-dorsolateral gradient, separating the limbic and paralimibic systems from posterior parietal cortex. Another pattern, characterized by dynamic range, follows a unimodal-transmodal gradient, separating primary sensory-motor cortices from association cortex. Importantly, the two patterns are related to gradients of functional activation. The ventromedial-dorsolateral pattern differentiates affective versus cognitive activation, whereas the unimodal-transmodal pattern differentiates primary sensory versus higher-order cognitive processing. Collectively, these results provide evidence that local computations reflect systematic variation in multiple anatomical circuit properties, ultimately manifesting as unique temporal signatures in regional activity and patterns of functional specialization.

An emerging literature emphasizes the hierarchical organization of neural systems, whereby systematic variations in laminar architecture across the cortical sheet are mirrored by multiple cytological properties, including neuron density, spine count, branching and neurotransmitter receptor profiles [47, 63, 65]. These variations ultimately manifest as spatially ordered gradients of structural and functional attributes [50], including gene expression [12, 31], cortical thickness [100], intra-cortical myelin [49], laminar differentiation [72, 99] and excitability [22, 64, 88, 102]. Indeed, we find that the two patterns of intrinsic dynamics are closely related to gene expression, intracortical myelin and cortical thickness. Our results build on this literature, demonstrating that microscale and connectional hierarchies leave an indelible mark on intrinsic dynamics [59], perhaps through variation in local excitability [102]. How these patterns are related to underlying cell types and subcortical afferent input – in particular, thalamocortical feedback – is an important ongoing question [1, 34, 70, 73, 85, 101].

More generally, the present findings are part of a larger trend in the field to understand structure-function relationships by considering molecular [4, 28, 77, 107], cellular [3, 70, 81, 83] and physiological [25, 82] attributes of network nodes, thereby conceptually linking local and global brain organization [55, 89]. In such “annotated networks”, macroscale network architecture is thought to reflect similarity in local properties, and *vice versa*, such that areas with similar properties are more likely to be anatomically connected and to functionally interact with one another [11, 42, 46, 104]. Indeed, we find that two regions are more likely to display similar intrinsic dynamics if they are anatomically connected and if they are part of the same functional community, suggesting that network organization and local intrinsic dynamics are intertwined. A significant corollary of the present work is that functional connectivity – presently conceptualized as coherent fluctuations in neural activity and operationalized as correlated BOLD values over time – misses out on an important set of inter-regional relationships. Namely, two regions may display identical dynamic profiles, suggesting common function, but unless they also display time-locked activity, current methods would miss out on this potentially biologically meaningful inter-regional relationship.

The present results are consistent with contemporary theories linking brain structure and function, but they must be interpreted with respect to several methodological caveats. First, all present analyses were performed on BOLD time series with lower sampling rate compared to electromagnetic recordings, potentially obscuring more subtle dynamics occurring on faster timescales. To mitigate this concern, all analyses were performed in high-resolution multiband HCP data with multiple runs, and replicated in MSC data, but in principle, the present analyses could be repeated and validated in magnetoencephalographic recordings. Second, all analyses were performed on haemodynamic time courses that may not completely reflect the underlying neuronal population dynamics. Despite this caveat, we observe a close correspondence between the isolated patterns of intrinsic dynamics and molecular, structural, functional, and psychological gradients. Third, the pattern of temporal signal-to-noise ratio in the BOLD is known to be nonuniform, but it is not correlated with the intrinsic dynamics patterns observed in the present report.

Altogether, the present results point towards highly patterned intrinsic dynamics across the neocortex. These patterns reflect prominent molecular and microstructural gradients, as well as macroscale structural and functional organization. Importantly, patterns of intrinsic dynamics drive spatial variation in functional activation. These findings demonstrate that structural organization of the brain shapes patterns of intrinsic dynamics, ultimately manifesting as distinct axes of psychological processes.

## METHODS

### Dataset: Human Connectome Project (HCP)

Following the procedure described in [21], we obtained structural and functional magnetic resonance imaging (MRI) data of two sets of healthy young adults (age range 22-35 years) with no familial relationships (neither within nor between sets) as *Discovery* (*n* = 201) and *Validation* (*n* = 127) sets from Human Connectome Project (HCP; S900 release [93]). All four resting state fMRI scans (2 scans (R/L and L/R phase encoding directions) on day one and 2 scans (R/L and L/R phase encoding directions) on day two, each about 15 minutes long; TR = 720 ms), as well as structural MRI and diffusion weighted imaging (DWI) data were available for all participants.

### HCP Data Processing

All the structural and functional MRI data were preprocessed using HCP minimal pre-processing pipelines [38, 93]. We provide a brief description of data preprocessing below, while detailed information regarding data acquisition and pre-processing is available elsewhere [38, 93]. The procedure was separately repeated for *Discovery* and *Validation* sets.

### Structural MRI

T1- and T2-weighted MR images were corrected for gradient nonlinearity, and when available, the images were co-registered and averaged across repeated scans for each individual. The corrected T1w and T2w images were co-registered and cortical surfaces were extracted using FreeSurfer 5.3.0-HCP [20, 21, 26]. For each individual, cortical thickness was estimated as the difference between pial and white matter surfaces and intracortical myelin content was estimated as T1w/T2w ratio. The pre-processed data were parcellated into 400 cortical areas using Schaefer parcellation [80].

### Resting state functional MRI

All 3T functional MRI time series were corrected for gradient nonlinearity, head motion using a rigid body transformation, and geometric distortions using scan pairs with opposite phase encoding directions (R/L, L/R) [21]. Further pre-processing steps include coregistration of the corrected images to the T1w structural MR images, brain extraction, normalization of whole brain intensity, high-pass filtering (> 2000s FWHM; to correct for scanner drifts), and removing additional noise using the ICA-FIX process [21, 79]. The pre-processed time series were then parcellated into 400 areas as described above. The parcellated time series were used to construct functional connectivity matrices as a Pearson correlation coefficient between pairs of regional time series for each of the four scans of each participant. A group-average functional connectivity matrix was constructed as the mean functional connectivity across all individuals and scans.

### Diffusion weighted imaging (DWI)

DWI data was pre-processed using the MRtrix3 package [91] (https://www.mrtrix.org/). More specifically, fiber orientation distributions were generated using the multi-shell multi-tissue constrained spherical deconvolution algorithm from MRtrix [24, 52]. White matter edges were then reconstructed using probabilistic streamline tractography based on the generated fiber orientation distributions [90]. The tract weights were then optimized by estimating an appropriate cross-section multiplier for each streamline following the procedure proposed by Smith and colleagues [86] and a connectivity matrix was built for each participant using the same parcellation as described above. Finally, we used a consensus approach to construct a binary group-level structural connectivity matrix, preserving the edge length distribution in individual participants [10, 67, 68, 83].

### Replication dataset: Midnight Scan Club (MSC)

We used resting state fMRI data of *n* = 10 healthy young adults, each with 10 scan sessions of about 30 minutes long, from Midnight Scan Club (MSC [41]) dataset as an independent replication dataset. Details about the participants, MRI acquisition, and data pre-processing are provided by Gordon and colleagues elsewhere [41]. We obtained the surface-based, preprocessed resting state fMRI time courses in CIFTI format through OpenNeuro (https://openneuro.org/datasets/ds000224/versions/1.0.0). The pre-processing steps include motion correction and global signal regression [41]. Following the pre-processing methods suggested by Gordon and colleagues [41], we smoothed the surface-level time series data with geodesic 2D Gaussian kernels (σ = 2.55mm) using the Connectome Workbench [62]. Finally, we censored the motion-contaminated frames of time series for each participant separately, using the temporal masks provided with the dataset. The pre-processed data were parcellated into 400 cortical regions using Schaefer parcellation [80]. One participant (MSC08) was excluded from subsequent analysis due to low data reliability and self-reported sleep as described in [41]. The parcellated time series were then subjected to the same analyses that were performed on the HCP *Discovery* and *Validation* datasets.

### Microarray expression data: Allen Human Brain Atlas (AHBA)

Regional microarray expression data were obtained from six post-mortem brains provided by the Allen Human Brain Atlas (AHBA; http://human.brain-map.org/) [45]. We used the *abagen* (https://github.com/netneurolab/abagen) toolbox to process and map the data to 400 parcellated brain regions from Schaefer parcellation [80].

Briefly, genetic probes were reannotated using information provided by [5] instead of the default probe information from the AHBA dataset. Using reannotated information discards probes that cannot be reliably matched to genes. The reannotated probes were filtered based on their intensity relative to background noise levels [76]; probes with intensity less than background in ≥50% of samples were discarded. A single probe with the highest differential stability, Δ_*S*_ (*p*), was selected to represent each gene [44], where differential stability was calculated as:

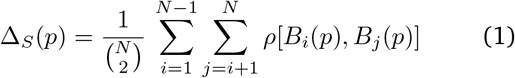

Here, *ρ* is Spearman’s rank correlation of the expression of a single probe *p* across regions in two donor brains, *B_i_* and *B_j_*, and *N* is the total number of donor brains. This procedure retained 15,656 probes, each representing a unique gene.

Next, tissue samples were mirrored across left and right hemispheres [78] and then assigned to brain regions using their corrected MNI coordinates (https://github.com/chrisfilo/alleninf) by finding the nearest region, up to 2mm away. To reduce the potential for misassignment, sample-to-region matching was constrained by hemisphere and cortical/subcortical divisions [5]. If a brain region was not assigned any sample based on the above procedure, the sample closest to the centroid of that region was selected in order to ensure that all brain regions were assigned a value. Samples assigned to the same brain region were averaged separately for each donor. Gene expression values were then normalized separately for each donor across regions using a robust sigmoid function and rescaled to the unit interval [28]. Scaled expression profiles were finally averaged across donors, resulting in a single matrix with rows corresponding to brain regions and columns corresponding to the retained 15,656 genes. The expression values of 1,906 brain-specific genes were used for further analysis [12].

### Massive temporal feature extraction using *hctsa*

We used the highly comparative time series analysis toolbox, *hctsa* [29, 30], to perform a massive feature extraction of the time series of each brain area for each participant. The *hctsa* package extracted over 7,000 local temporal properties using a wide range of operations based on time series analysis [29, 30]. The extracted features include, but are not limited to, distributional properties, entropy and variability, autocorrelation, timedelay embeddings, and nonlinear properties of a given time series.

The *hctsa* feature extraction analysis was performed on the parcellated fMRI time series of each run and each participant separately (Fig. 1). Following the feature extraction procedure, the outputs of the operations that produced errors were removed and the remaining features (above 6,000 features) were normalized across nodes using an outlier-robust sigmoidal transform. We used Pearson correlation coefficients to measure the pairwise similarity between the temporal features of all possible combinations of brain areas. As a result, a temporal profile similarity network was constructed for each individual and each run, representing the strength of the similarity of the local temporal fingerprints of brain areas (Fig. 1). The resulting similarity matrices were then compared to the underlying functional and structural brain networks.

### Neurosynth

Functional activation probability maps were obtained for multiple psychological terms using Neurosynth [105] (https://github.com/neurosynth/neurosynth). Probability maps were restricted to those for terms present in both Neurosynth and the Cognitive Atlas [74], yielding a total of 123 maps (Table S1). We used the volumetric “association test” (i.e. reverse inference) maps, which were projected to the FreeSurfer *fsaverage5* mid-grey surface with nearest neighbor interpolation using Freesurfer’s *mri_vol2surf* function (v6.0.0; http://surfer.nmr.mgh.harvard.edu/). The resulting surface maps were then parcellated to 400 cortical regions using the Schaefer parcellation [80].

### Null model

A consistent question in the present work is the topographic correlation between temporal features and other features of interest. To make inferences about these links, we implement a null model that systematically disrupts the relationship between two topographic maps but preserves their spatial autocorrelation [2] (see also [12, 13] for an alternative approach). We first created a surface-based representation of the Cammoun atlas on the FreeSurfer fsaverage surface using the Connectome Mapper toolkit (https://github.com/LTS5/cmp; [19]). We used the spherical projection of the *fsaverage* surface to define spatial coordinates for each parcel by selecting the vertex closest to the center-of-mass of each parcel [83, 94, 95]. The resulting spatial coordinates were used to generate null models by applying randomly-sampled rotations and reassigning node values based on the closest resulting parcel (10,000 repetitions). The rotation was applied to one hemisphere and then mirrored to the other hemisphere.

## ACKNOWLEDGMENTS

We thank Vincent Bazinet, Justine Hansen, Estefany Suarez, Bertha Vazquez-Rodriguez and Zhen-Qi Liu for helpful comments and stimulating discussion. This research was undertaken thanks in part to funding from the Canada First Research Excellence Fund, awarded to McGill University for the Healthy Brains for Healthy Lives initiative. BM acknowledges support from the Natural Sciences and Engineering Research Council of Canada (NSERC Discovery Grant RGPIN #017-04265) and from the Canada Research Chairs Program. GS acknowledges support from the Natural Sciences and Engineering Research Council of Canada (NSERC).

---

**TABLE S1.** List of terms used in Neurosynth analyses. The overlapping terms between Neurosynth [105] and Cognitive Atlas [74] corpuses used in the reported analyses are listed below.

|  |  |  |  |  |
| --- | --- | --- | --- | --- |
| action | eating | insight | naming | semantic memory |
| adaptation | efficiency | integration | navigation | sentence comprehension |
| addiction | effort | intelligence | object recognition | skill |
| anticipation | emotion | intention | pain | sleep |
| anxiety | emotion regulation | interference | perception | social cognition |
| arousal | empathy | judgment | planning | spatial attention |
| association | encoding | knowledge | priming | speech perception |
| attention | episodic memory | language | psychosis | speech production |
| autobiographical memory | expectancy | language comprehension | reading | strategy |
| balance | expertise | learning | reasoning | strength |
| belief | extinction | listening | recall | stress |
| categorization | face recognition | localization | recognition | sustained attention |
| cognitive control | facial expression | loss | rehearsal | task difficulty |
| communication | familiarity | maintenance | reinforcement learning | thought |
| competition | fear | manipulation | response inhibition | uncertainty |
| concept | fixation | meaning | response selection | updating |
| consciousness | focus | memory | retention | utility |
| consolidation | gaze | memory retrieval | retrieval | valence |
| context | goal | mental imagery | reward anticipation | verbal fluency |
| coordination | hyperactivity | monitoring | rhythm | visual attention |
| decision | imagery | mood | risk | visual perception |
| decision making | impulsivity | morphology | rule | word recognition |
| detection | induction | motor control | salience | working memory |
| discrimination | inference | movement | search |  |
| distraction | inhibition | multisensory | selective attention |  |

**Figure S1.**
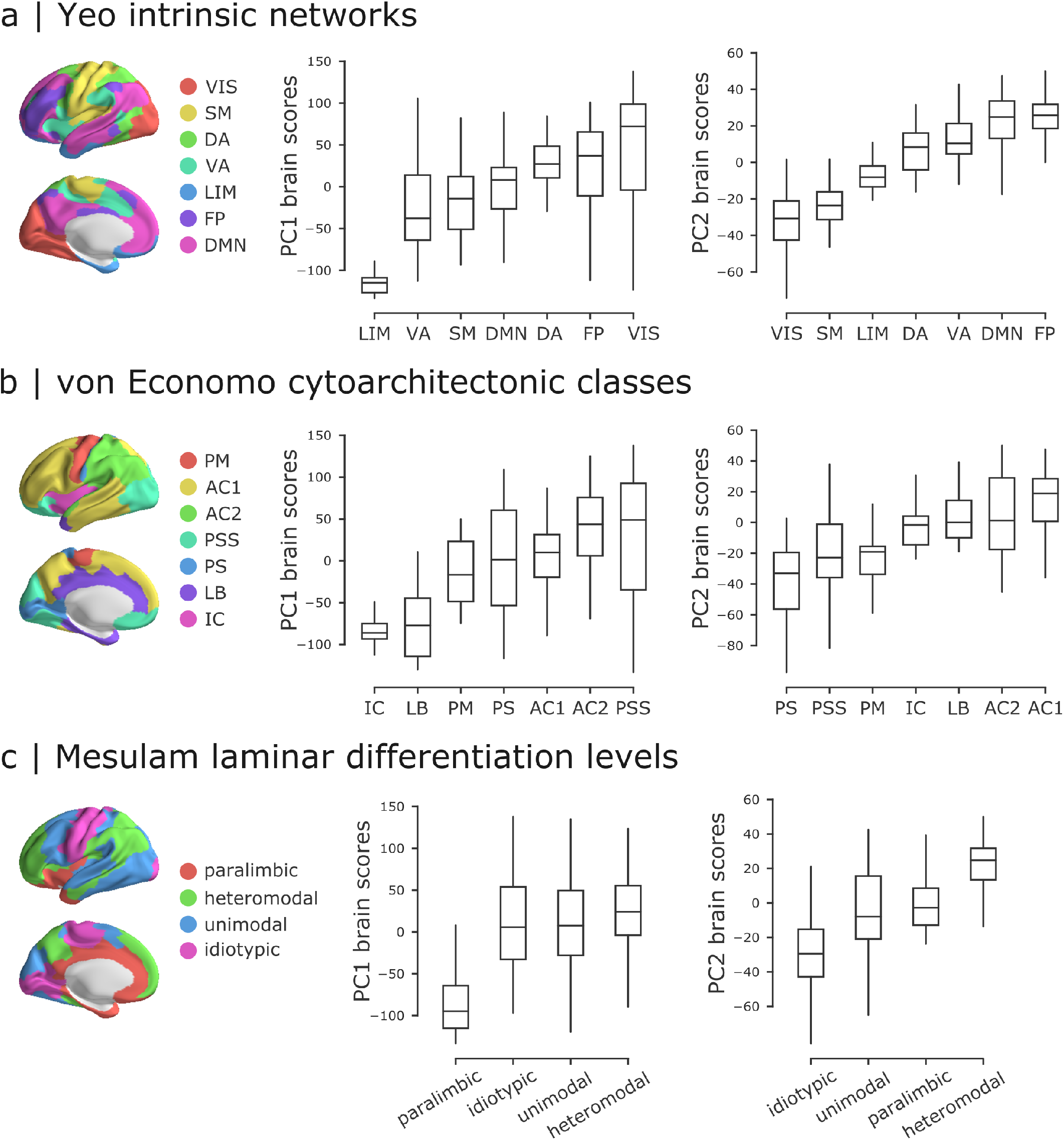
Intrinsic dynamics across intrinsic networks, cytoarchitectonic classes and laminar differentiation levels. Mean PC1 and PC2 scores were computed for the constituent classes in three commonly used anatomical and functional partitions of the brain: (a) intrinsic fMRI networks [106], (b) cytoarchitectonic classes [96–98], laminar differentiation levels [66]. Intrinsic networks: VIS = visual, SM = somatomotor, DA = dorsal attention, VA = ventral attention, LIM = limbic, FP = fronto-parietal, DMN = default mode. Cytoarchitectonic classes: PM = primary motor cortex, AC1 = association cortex, AC2 = association cortex, PSS = primary/secondary sensory, PS = primary sensory cortex, LB = limbic regions, IC = insular cortex.

